# Genetic compensation is triggered by mutant mRNA degradation

**DOI:** 10.1101/328153

**Authors:** Mohamed A. El-Brolosy, Andrea Rossi, Zacharias Kontarakis, Carsten Kuenne, Stefan Günther, Nana Fukuda, Carter Takacs, Shih-Lei Lai, Ryuichi Fukuda, Claudia Gerri, Khrievono Kikhi, Antonio J. Giraldez, Didier Y.R. Stainier

## Abstract

Genetic compensation by transcriptional modulation of related gene(s) (also known as transcriptional adaptation) has been reported in numerous systems ^1–3^; however, whether and how such a response can be activated in the absence of protein feedback loops is unknown. Here, we develop and analyze several models of transcriptional adaptation in zebrafish and mouse that we show are not caused by loss of protein function. We find that the increase in transcript levels is due to enhanced transcription, and observe a correlation between the levels of mutant mRNA decay and transcriptional upregulation of related genes. To assess the role of mutant mRNA degradation in triggering transcriptional adaptation, we use genetic and pharmacological approaches and find that mRNA degradation is indeed required for this process. Notably, uncapped RNAs, themselves subjected to rapid degradation, can also induce transcriptional adaptation. Next, we generate alleles that fail to transcribe the mutated gene and find that they do not show transcriptional adaptation, and exhibit more severe phenotypes than those observed in alleles displaying mutant mRNA decay. Transcriptome analysis of these different alleles reveals the upregulation of hundreds of genes with enrichment for those showing sequence similarity with the mutated gene’s mRNA, suggesting a model whereby mRNA degradation products induce the response via sequence similarity. These results expand the role of the mRNA surveillance machinery in buffering against mutations by triggering the transcriptional upregulation of related genes. Besides implications for our understanding of disease-causing mutations, our findings will help design mutant alleles with minimal transcriptional adaptation-derived compensation.

---

Recent advances in reverse genetic tools have greatly enhanced our ability to study gene function in a much wider range of organisms. These studies have reinforced previous observations that many engineered mutants do not exhibit an obvious phenotype, reviving interest in the concept of genetic robustness. Several mechanisms have been proposed to explain genetic robustness, including functional redundancy^4^, rewiring of genetic networks^5^, and the acquisition of adaptive mutations in the case of rapidly proliferating organisms such as yeast^6^. In a previous report^1^, we proposed genetic compensation as another underlying mechanism, whereby a deleterious mutation can lead to the transcriptional upregulation of related genes which can assume the function of the mutated gene. We provided evidence that this upregulation is induced upstream of the loss of protein function, implying the existence of an unknown trigger. In order to investigate the molecular machinery underlying genetic compensation, we developed and investigated zebrafish and mouse mutants that display transcriptional adaptation, a form of genetic compensation that involves the transcriptional upregulation of genes that can potentially compensate for the loss of the mutated gene.

We started by analyzing different zebrafish mutants, namely *hbegfa*, *vcla*, *hif1ab*, *vegfaa*, *egfl7* and *alcama* mutants, and found that they transcriptionally upregulate a paralogue or a family member (hereafter referred to as ‘adapting genes’), namely *hbegfb*, *vclb*, *epasla* and *epaslb*, *vegfab*, *emilin3a* and *alcamb*, respectively (Fig. 1a). Moreover, we found that *vcla*, *hif1ab* and *egfl7* heterozygous animals also display a transcriptional adaptation response, albeit less pronounced than that observed in the homozygous mutants (Extended Data Fig. 1a), indicating that transcriptional adaptation is a dominant phenomenon. We also observed upregulation of the *hbegfa*, *hif1ab*, *vegfaa* and *alcama* wild-type (wt) alleles in the respective heterozygous mutants (Extended Data Fig. 1b). Injection of wt *hif1ab*, *vegfaa*, *egfl7* and *alcama* mRNA into the respective mutants did not dampen the transcriptional adaptation response (Fig. 1b), indicating that the response is triggered upstream of the loss of protein function. These data are consistent with our previous observations that embryos injected with a dominant negative version of Vegfaa do not exhibit upregulation of *vegfab* mRNA levels even though they appear similar to *vegfaa* mutants^1^. Similarly, we found that *Kindlin-2* mutant mouse kidney fibroblasts (MKFs), *Rela* and *Actg1* mutant mouse embryonic fibroblasts (MEFs) and *Actb* mutant mouse embryonic stem cells (mESCs) (hereafter referred to as the knockout (K.O.) alleles) transcriptionally upregulate *Kindlin-1*, *Rel*, *Actg2* and *Actg1*, respectively (Fig. 1c). *Actb* heterozygous mESCs also upregulate *Actg1* (Extended Data Fig 1b). Transfection of wt *Kindlin-2* and *Rela* in the respective mutant cells also did not dampen the transcriptional adaptation response (Fig. 1d and Extended Data Fig. 2). Altogether, these data strongly indicate that the loss of protein function is not the trigger for the transcriptional adaptation response observed in these models.

**Figure 1.**
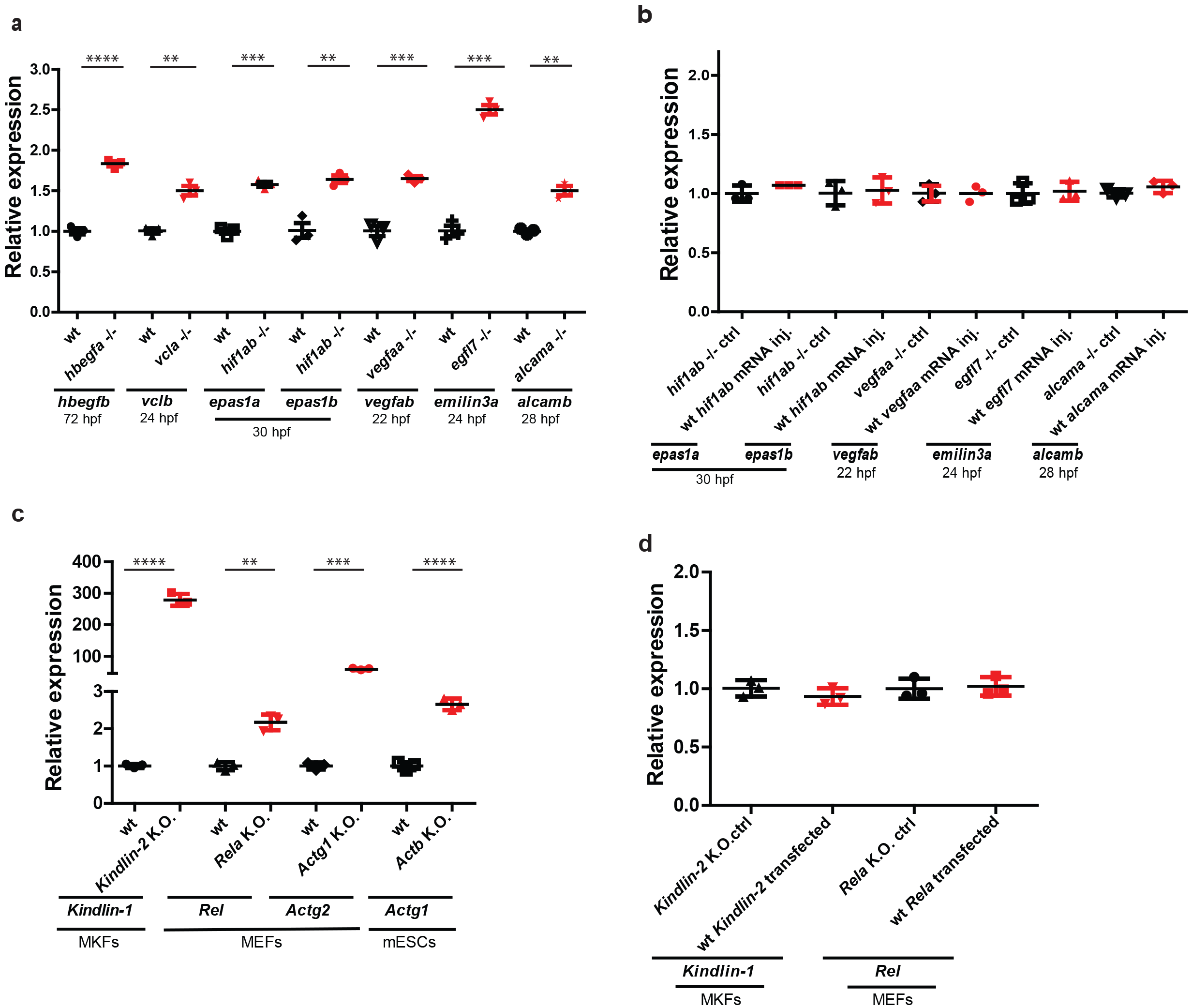
Transcriptional adaptation models in zebrafish and mouse. **a**, qPCR analysis of *hbegfb*, *vclb*, *epasla* and *epaslb*, *vegfab*, *emilin3a* and *alcamb* mRNA levels in *hbegfa*, *vcla*, *hiflab*, *vegfaa*, *egfl7* and *alcama* wt and mutant zebrafish. **b**, qPCR analysis of *epasla* and *epaslb*, *vegfab*, *emilin3a* and *alcamb* mRNA levels at 24 hours post fertilization (hpf) in *hiflab*, *vegfaa*, *egfl7* and *alcama* mutant embryos injected with *eGFP* mRNA (control) or wt mRNA. **c**, qPCR analysis of *Kindlin-1*, *Rel*, *Actg2* and *Actgl* mRNA levels in *Kindlin-2*, *Rela*, *Actgl* and *Actb* wt and knockout cells. **d**, qPCR analysis of *Kindlin-1* and *Rel* mRNA levels in *Kindlin-2* and *Rela* knockout cells transfected with empty vectors (control) or plasmids encoding wt KINDLIN-2 or RELA. **a-d**, Data points represent different biological replicates. Wt or control expression set at 1 for each assay. Error bars, s.d. Two-tailed student’s *t*-test was used to assess *P* values. \*\**P* ≤ 0.01, \*\*\**P* ≤ 0.001, **** *P* ≤ 0.0001.

To determine whether the increase in mRNA levels was due to increased transcription of the adapting genes or increased mRNA stability, we measured pre-mRNA levels of *hbegfb* and *emilin3a* in *hbegfa* and *egfl7* mutants and found that they were upregulated to a similar extent as the mature mRNA (Extended Data Fig. 3a). Similar findings were obtained for *Kindlin-1* and *Rel* pre-mRNA levels in *Kindlin-2* and *Rela* K.O. cells (Extended Data Fig. 3b). Together, these data imply an increase in transcription of the adapting genes. To investigate the chromatin changes accompanying the transcriptional adaptation response, we performed ATAC-seq and found that *Kindlin-2* K.O. MEFs display increased chromatin accessibility at the *Kindlin-1* promoter region as well as at a potential enhancer region recently identified by Hi-C analysis ^7, 8^ (Extended Data Fig. 3c). Similarly, the *Rela* K.O. MEFs display increased chromatin accessibility at a potential *Rel* enhancer region, also identified by Hi-C analysis ^7, 8^ (Extended Data Fig. 3d).

Since the loss of protein function does not appear to be the trigger for the transcriptional adaptation response, we investigated two other possible triggers, the DNA lesion and the mutant mRNA. We first reasoned that if the DNA lesion itself was the trigger for the transcriptional adaptation response, all mutations should induce it. However, some *hbegfa*, *vcla*, *vegfaa* and *egfl7* mutant alleles do not display transcriptional adaptation (Extended Data Fig. 4a, b, c), indicating that the DNA lesion is not the trigger. While analyzing various mutant alleles, we found that two different alleles for *hbegfa* and *vcla* behave differently in terms of transcriptional adaptation. While the previously mentioned CRISPR-generated alleles of *hbegfa* (hereafter referred to as *hbegfa*^*Δ7*^) and *vcla* (hereafter referred to as *vcla*^*Δ13*^) display transcriptional upregulation of *hbegfb* and *vclb*, respectively, ENU-induced alleles (*hbegfa*^*sa18135*^ and *vcla*^*sa14599*^) do not display transcriptional adaptation (Fig. 2a). For both genes, both alleles lead to a premature termination codon (PTC) in similar regions of the coding sequence. To investigate these differences further, we examined the degradation of the mutant mRNA in the various alleles, and observed that the extent of the transcriptional adaptation response correlated with the levels of mutant mRNA decay. For example, while the *hbegfa*^*Δ7*^ allele displays a 50% reduction in mutant transcript levels, the *hbegfa*^*sa18135*^ allele displays only a 20% decrease; and similarly, while the *vcla*^*Δ13*^allele displays an 80% reduction in mutant transcript levels, the *vcla*^*sa14599*^ allele displays no noticeable decrease (Fig. 2b). Moreover, we observed that all of the previously mentioned zebrafish and mouse models of transcriptional adaptation display mutant mRNA decay (Fig. 2c, d). In contrast, alleles that fail to induce a transcriptional adaptation response do not display mutant mRNA decay (Extended Data Fig. 4a, b, c). To confirm that the observed reduction of mutant transcript levels was due to mRNA decay and not decreased transcription of the mutated gene, we analyzed the pre-mRNA levels of *hbegfa*, *egfl7* and *alcama* in *hbegfa*^*Δ7*^, *egfl7* and *alcama* mutant zebrafish, and found that unlike the mRNA, the pre-mRNA levels remained unchanged, or were slightly upregulated, compared to wt (Extended Data Fig. 5a). Similar findings were observed in *Kindlin-2* and *Rela* K.O. cells (Extended Data Fig. 5b). Thus, mutants that exhibit transcriptional adaptation show reduced mutant mRNA levels due to mRNA decay.

**Figure 2.**
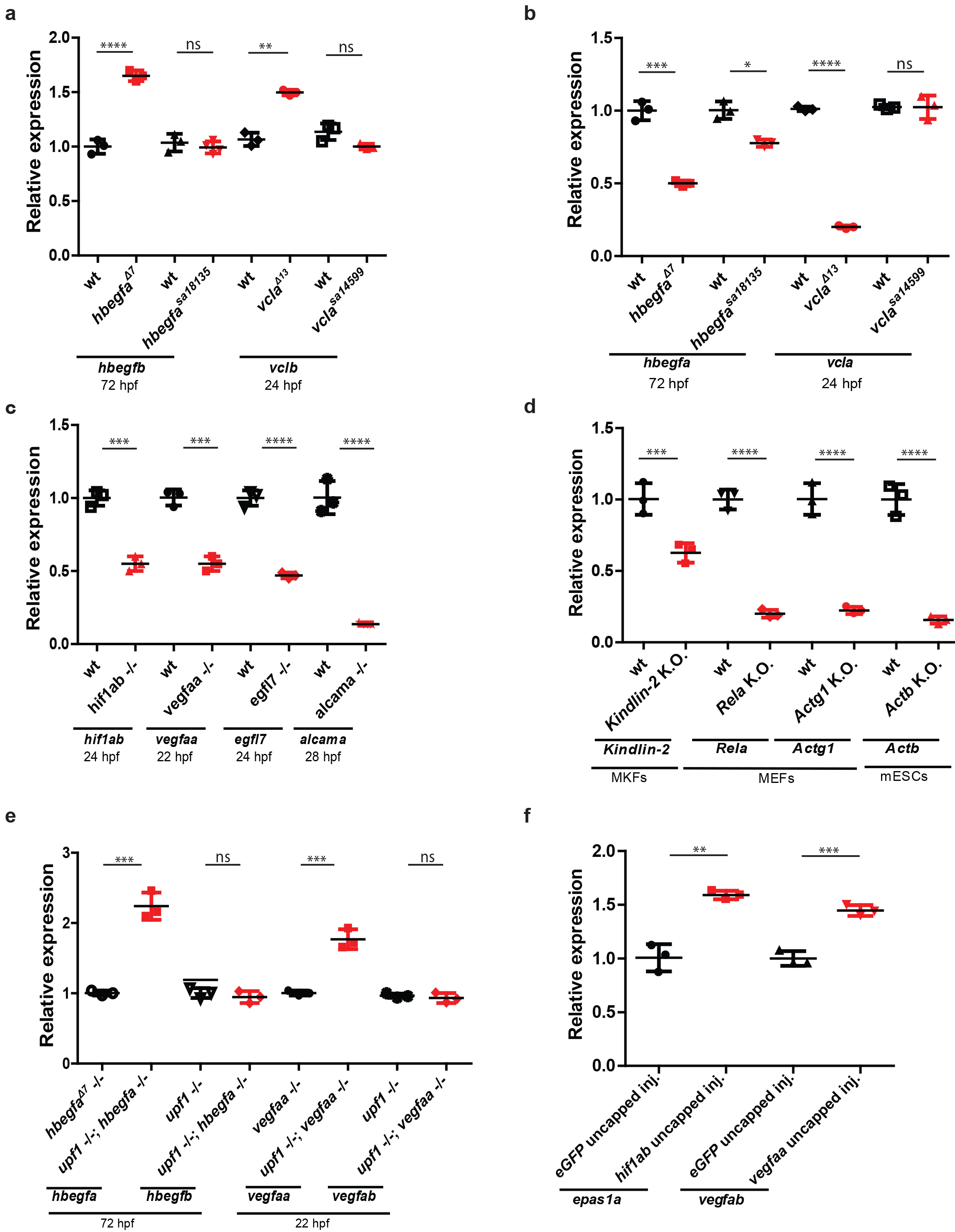
Mutant mRNA decay induces transcriptional adaptation. **a**, qPCR analysis of *hbegfb* and *vclb* mRNA levels in the indicated *hbegfa* and *vcla* mutant alleles. **b**, qPCR analysis of *hbegfa* and *vcla* mRNA levels in the indicated *hbegfa* and *vcla* mutant alleles. **c**, qPCR analysis of *hiflab*, *vegfaa*, *egfl7* and *alcama* mRNA levels in *hiflab*, *vegfaa*, *egfl7* and *alcama* wt and mutant zebrafish. **d**, qPCR analysis of *Kindlin-2*, *Rela*, *Actgl* and *Actb* mRNA levels in *Kindlin-2*, *Rela*, *Actgl* and *Actb* wt and knockout cells. **e**, qPCR analysis of *hbegfa*, *hbegfb* and *vegfaa*, *vegfab* mRNA levels in *upfl;hbegfa* or *upfl;vegfaa* double mutant zebrafish compared to the indicated controls. **f**, qPCR analysis of *epas1a* and *vegfab* mRNA levels in 6 hpf wt embryos injected with uncapped *hiflab* and *vegfaa* mRNA. **a-f**, Data points represent different biological replicates. Wt or control expression set at 1 for each assay. Error bars, s.d. Two-tailed student’s *t*-test was used to assess *P* values. \**P* < 0.05, \*\**P* ≤ 0.01, \*\*\**P* ≤ 0.001, **** *P* ≤ 0.0001, ns: not significant.

Mutant transcripts with a PTC are usually degraded through the nonsense-mediated decay (NMD) pathway. Upf1 is one of the key proteins that help detect PTCs and trigger the NMD pathway^9^. To investigate the role of the mRNA surveillance machinery in triggering transcriptional adaptation to mutations, we genetically inactivated the NMD pathway in *hbegfa*^*Δ7*^, *vegfaa* and *vcla*^*Δ13*^ zebrafish mutants. Inactivating *upf1* in these mutants led to decreased levels of mutant mRNA decay and also to the loss of transcriptional adaptation (Fig. 2e and Extended Data Fig. 6a). Similar findings were obtained when knocking down UPF1 and EXOSC4 (a component of the exosome complex required for 3’ to 5’ degradation of defective transcripts) in *Rela* K.O. MEFs, or when knocking down SMG6 (another key protein of the mRNA surveillance machinery) in *Actb* K.O. mESCs (Extended Data Fig. 6b, c), and also when pharmacologically inhibiting NMD in *hbegfa*^*Δ7*^ zebrafish mutants (Extended Data Fig. 6d).

We next asked whether inducing mRNA decay in wt zebrafish or mouse cells by using uncapped mRNAs, which are known to be rapidly degraded by 5’ to 3’ exonucleases^10^, would induce a transcriptional adaptation response. Indeed, we found that injection of uncapped *hif1ab* or *vegfaa* mRNAs into wt embryos induced a transcriptional adaptation response (Fig. 2f and Extended Data Fig. 6e) as well as an increase in endogenous *hif1ab* and *vegfaa* gene expression (data not shown). Similarly, transfection of uncapped *Actb* into wt mESCs led to *Actg1* upregulation (Extended Data Fig. 6f). Notably, injection of uncapped *hif1ab* or *vegfaa* mRNAs with an upstream sequence which makes them resistant to 5’ to 3’ exonuclease-mediated decay^11^ did not induce a transcriptional adaptation response (data not shown). Taken together, these data indicate that mRNA degradation is sufficient to induce a transcriptional adaptation response.

Next, we reasoned that if the process of mutant mRNA degradation is the trigger for the transcriptional adaptation response, alleles that fail to transcribe the mutated gene should not display this response. To this end, we used CRISPR/Cas9 technology to generate such alleles (hereafter referred to as RNA-less alleles) through deletion of the promoter region or the entire gene locus (for compact genes). Indeed, RNA-less alleles of *hbegfa*, *vegfaa* and *alcama* failed to upregulate *hbegfb*, *vegfab* and *alcamb* (Fig. 3a). Similarly, in mouse cells, RNA-less alleles of *Rela, Actg1* and *Actb* failed to exhibit a transcriptional adaptation response (Fig. 3b). We also attempted to generate a promoter-less allele for *Kindlin-2* in MKFs; however, the obtained clones exhibited proliferation defects that prevented their expansion. As an alternative, we used CRISPR interference (CRISPRi) and found that reducing transcription of the mutant *Kindlin-2* gene in *Kindlin-2* K.O. cells led to a decrease in *Kindlin-1* transcriptional upregulation (Extended Data Fig. 6g). RELA and REL are NF-κB transcription factors that play an essential role in preventing tumor necrosis factor α (TNFα) mediated apoptosis^12^. We observed that the promoter-less *Rela* MEFs were more sensitive to TNFα-induced apoptosis than the *Rela* K.O. MEFs (Fig. 3c). Moreover, mESCs with an *Actb* full locus deletion displayed less protrusive activity and more severe growth defects than *Actb* K.O. cells did (Fig. 3d, e). We also generated an RNA-less allele for *egfl7* and observed that the mutant embryos display pronounced vascular defects (Fig. 3f). Therefore, generation of RNA-less alleles can uncover phenotypes masked by transcriptional adaptation-derived compensation in mutant alleles that display mRNA decay.

**Figure 3.**
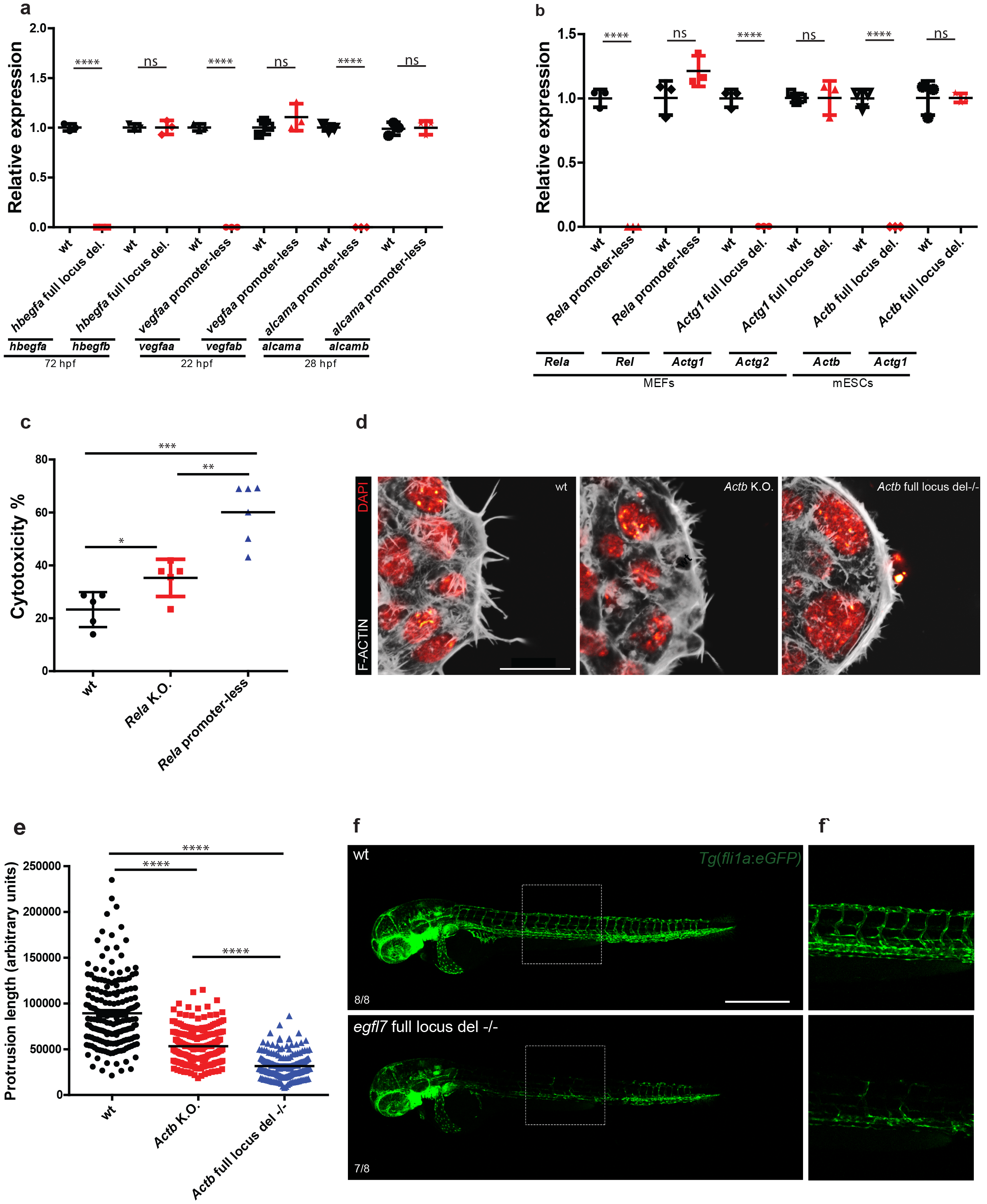
Alleles that fail to transcribe the mutated gene do not display transcriptional adaptation and exhibit stronger phenotypes. **a**, qPCR analysis of *hbegfa*, *hbegfb*, *vegfaa*, vegfab, alcama and *alcamb* mRNA levels in zebrafish lacking the full *hbegfa* locus or the *vegfaa* or *alcama* promoter compared to wt siblings. **b**, qPCR analysis of *Rela*, *Rel*, *Actb*, *Actgl*, *Actgl* and *Actg2* mRNA levels in MEFs and mESCs lacking the *Rela* promoter or the full locus of *Actg1* or *Actb* compared to wt cells. **c**, Cytotoxicity assay 24 hours following treatment of wt, *Rela* K.O. and *Rela* promoter-less MEFs with 25 ng/ml mouse TNFα. Percentages were normalized relative to DMSO-treated cells. **d**, Confocal micrographs of wt, *Actb* K.O. and *Actb* full locus deletion mESCs. Actin filaments are depicted in white and nuclei in red. **e**, Actin filament protrusion length (in arbitrary units) in wt, *Actb* K.O. and *Actb* full locus deletion mESCs. **f**, Confocal micrographs of 48 hpf *Tg*(*fli1a:eGFP*) wt and *egfl7* full locus deletion −/− sibling embryos in lateral views. Higher magnifications of dashed boxes are shown in **f’**. Scale bars: **e**: 20 μm, **f**: 500 μm. **a, b**, Wt or control expression set at 1 for each assay. **a-c, e**, Data points represent different biological replicates. Error bars, s.d. Twotailed student’s *t*-test was used to assess *P* values. \**P* ≤ 0.05, \*\**P* ≤ 0.01, \*\*\**P* ≤ 0.001, **** *P* ≤ 0.0001, ns: not significant.

Following mRNA decay, intermediates can be detected in the nucleus^13^, and small RNAs can modulate gene expression in a homology-mediated base-pairing fashion^14–17^. We thus reasoned that if mRNA decay intermediates act in a similar fashion to induce transcriptional adaptation, we should observe the upregulation of genes that exhibit sequence similarity with the mutated gene’s mRNA. Transcriptome analysis of *Kindlin-2*, *Actg1* and *Actb* K.O. cells revealed the upregulation of hundreds of genes in K.O. cells compared to wt, with no signs of a stress-induced response (Extended Data Fig. 7). Notably, we observed that at least 50% of protein-coding genes exhibiting sequence similarity with the mutated gene’s mRNA (hereafter referred to as ‘similar’ genes) were significantly upregulated, compared to a maximum of 21% when looking at a random set of protein-coding genes (Fig. 4a and Extended Data Fig. 8). In addition, 7 out of the 12 upregulated ‘similar’ genes in *Actg1* K.O. cells were not upregulated in *Actg1* RNA-less cells, and 4 out of the 6 upregulated ‘similar’ genes in *Actb* K.O. cells were not upregulated in *Actb* RNA-less cells (Extended Data Fig. 8). We then tested whether sequence similarity was sufficient to induce transcriptional adaptation. Mouse *Actb* and zebrafish *actb1* mRNAs exhibit a high degree of sequence similarity, and injection of uncapped mouse *Actb* mRNA into zebrafish embryos led to the upregulation of zebrafish *actb1* (Fig. 4b). On the other hand, injection of uncapped transcripts corresponding to the antisense strands of *hif1ab* and *vegfaa* mRNAs did not induce a transcriptional adaptation response (Fig. 4c), unlike what was observed with the sense strands (Fig. 2f). Moreover, injection of an uncapped synthetic transcript containing only sequences of *hif1ab* mRNA similar to the *epas1a* genomic locus was sufficient to induce a transcriptional adaptation response (Fig. 4d, Extended Data Fig. 9). Altogether these data suggest that transcriptional adaptation is induced in a sequence-similarity specific manner, possibly through mRNA decay intermediates (Fig. 4e).

**Figure 4.**
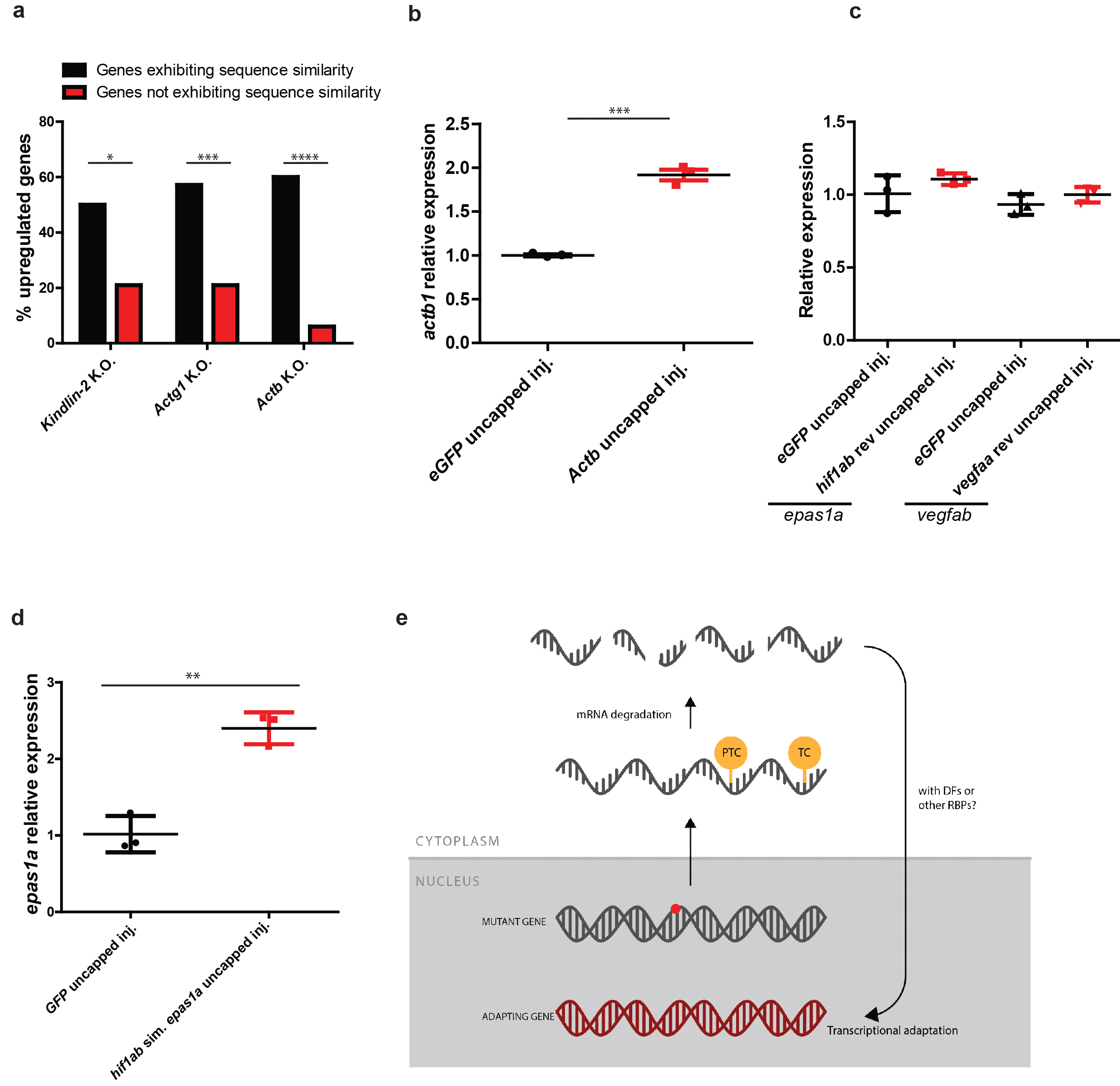
Upregulation of genes exhibiting sequence similarity. **a**, RNA-seq analysis showing percentage of significantly upregulated protein-coding genes (Log_2_ Fold Change K.O. > wt and *P* ≤ 0.05) for genes exhibiting sequence similarity with *Kindlin*-2, *Actg1* and *Actb* and for genes not exhibiting sequence similarity. **b**, qPCR analysis of *actb1* mRNA levels in 6 hpf wt embryos injected with uncapped mouse *Actb* mRNA. **c**, qPCR analysis of *epas1a* and *vegfab* mRNA levels in 6 hpf wt embryos injected with uncapped complementary transcripts of *hif1ab* and *vegfaa* mRNA. **d**, qPCR analysis of *epas1a* mRNA levels in 6 hpf wt embryos injected with uncapped RNA composed solely of the *hif1ab* sequences similar to *epas1a*. **e**, Current model of transcriptional adaptation to mutations. Red dot in mutated gene refers to a mutation. PTC: Premature termination codon; TC: Termination codon; DFs: Degradation factors; RBPs: RNA binding proteins. **b-d**, Control expression set at 1. Data points represent different biological replicates. Error bars, s.d. Two-tailed student’s *t*-test was used to assess *P* values. \**P* ≤ 0.05, \*\**P* ≤ 0.01, \*\*\**P* ≤ 0.001, **** *P* ≤ 0.0001.

Despite being a widely observed phenomenon^3^, transcriptional adaptation to mutations and its underlying molecular mechanisms remain poorly understood. Here we show that the mRNA surveillance machinery is important not only to prevent the translation of defective transcripts but also to buffer against mutations by triggering the transcriptional upregulation of related genes, including the wt allele of the mutated gene in the heterozygous state. In the past decade, it has become evident that the control of mRNA stability plays an important role in gene expression^18–21^. Previous studies have reported genetic and physical interactions of several mRNA decay factors with various proteins involved in gene expression, including RNA polymerase II and chromatin remodelers^22^. Indeed, mRNA decay factors can translocate to the nucleus and promote gene expression in a manner dependent on their ability to degrade mRNA^22^. Accordingly, we found that inactivating key mRNA decay factors leads to the loss of transcriptional adaptation (Fig. 2e and Extended Data Fig. 6b). Although mRNA decay intermediates were reported to be present in the nucleus^13^, their biological relevance remains unclear. A previous study reported that transfection of short fragments of the *Cdk9* or *Sox9* mRNA can lead to an increase in expression of these genes^15^. Mechanistically, these RNA fragments were found to downregulate antisense transcripts produced from these loci and which normally act as negative regulators of *Cdk9* and *Sox9* expression. Interestingly, we found that transfection of uncapped *Cdk9* or *Sox9* RNA leads to a clear upregulation of these genes (Extended Data Fig. 10a). Moreover, knockdown of a *BDNF* antisense transcript was reported to lead to the upregulation of the sense transcript, a response that involved a decrease in the negative histone mark H3K27me3^23^. Notably, we observed that transfection of uncapped *BDNF* sense transcript leads to a downregulation of the antisense transcript and a concomitant upregulation of the sense one (Extended Data Fig. 10b). These data indicate that acting on antisense transcripts is one possible mechanism through which mRNA decay intermediates can induce transcriptional adaptation in a sequence specific manner, a response that might also involve the modulation of histone marks.

Our study also provides clear guidelines towards generating mutant alleles that minimize compensation via transcriptional adaptation. We show that alleles that fail to transcribe the mutated gene do not exhibit transcriptional adaptation and can display phenotypes not observed in other mutant alleles (Fig. 3c-f). Consistent with our observations, a previous zebrafish study reported that mutant alleles for *mt2* with a lower degree of mutant mRNA decay display more severe phenotypes than alleles displaying a higher degree of mRNA decay^24^. For a number of human genetic diseases, several studies have reported that missense or in-frame indels, which are less likely to lead to mutant mRNA degradation, are more common in affected individuals than nonsense mutations or out-of-frame indels, which are more likely to lead to mutant mRNA degradation^25–30^. Interestingly, a study on Marfan syndrome patients reported that when compared to individuals with *FBN1* missense mutations, the mildest form of the disease was observed in an individual displaying very low mutant *FBNl* transcript levels due to an out-of-frame indel leading to a PTC in the *FBN1* coding sequence^31^. Similar results were observed in individuals with heterozygous nonsense mutations in the *HBB* gene; individuals who displayed decay of the mutant *HBB* transcripts were asymptomatic while individuals displaying no decay developed beta thalassemia-intermedia^32^. While the current dogma in the field is that missense mutations tend to be more common in some diseases as they may lead to constitutively active or dominant negative proteins, we propose that nonsense mutations might be less common in affected individuals as they might lead to mRNA decay-triggered compensatory upregulation of related genes and therefore not cause significant symptoms. Detailed transcriptomic analyses of relevant individuals will help test this hypothesis. Other studies have reported the upregulation of the wt allele in heterozygous conditions due to loss of negative feedback loops^33,34^. We found that transcriptional adaptation can also lead to the upregulation of the wt allele in heterozygous fish (Extended Data Fig. 1b), thus providing a possible explanation for haplosufficiency. Moreover, recent studies have reported homozygous loss-of-function mutations in several genes in healthy individuals (including genes such as *EGFL7* and *RELA* studied here)^35,36^. It will be interesting to investigate whether degradation of the mutant transcripts is associated with a transcriptional adaptation response that protects these individuals. Such analyses may help us understand why some mutations cause disease while others do not. They may also help identify new modifier genes whose expression levels could be further modulated for therapeutic purposes.

## Methods

Information on materials and methods can be found in the supplementary information.

## Acknowledgments

We thank Vahan Serobyan, Arica Beisaw and Felix Gunawan for comments on the manuscript and Jenny Pestel for the *alcama* mutant. Wt and *Kindlin-2* knockout MKFs were a generous gift from Reinhard Fässler (MPI for biochemistry, Martinsried, Germany). Wt and *Rela* knockout MEFs were a generous gift from Alexander Hoffmann (UCLA, USA). mESCs from the C57BL/6 mouse strain were a generous gift from Johnny Kim (MPI for heart and lung research, Bad Nauheim, Germany). We also thank Ann Atzberger for support with cell sorting and Martin Krzywinski for his help in drawing Figure 4e. M.A.E.B. was supported by a Boehringer Ingelheim Fonds PhD fellowship. Research in the Stainier lab is supported by the Max Planck Society, EU, DFG and Leducq Foundation.

## Author contributions

Performed experiments (M.A.E.B., A.R., Z.K., N.F., S.G., C.K., and K.K.), provided unpublished mutants (C.T., S.L., R.F. and C.G.), supervised experiments (A.J.G. and D.Y.R.S.) and wrote the manuscript (M.A.E.B. and D.Y.R.S.); all authors commented on the manuscript.

## Competing financial interests

The authors declare no competing financial interests.

## Figure legends

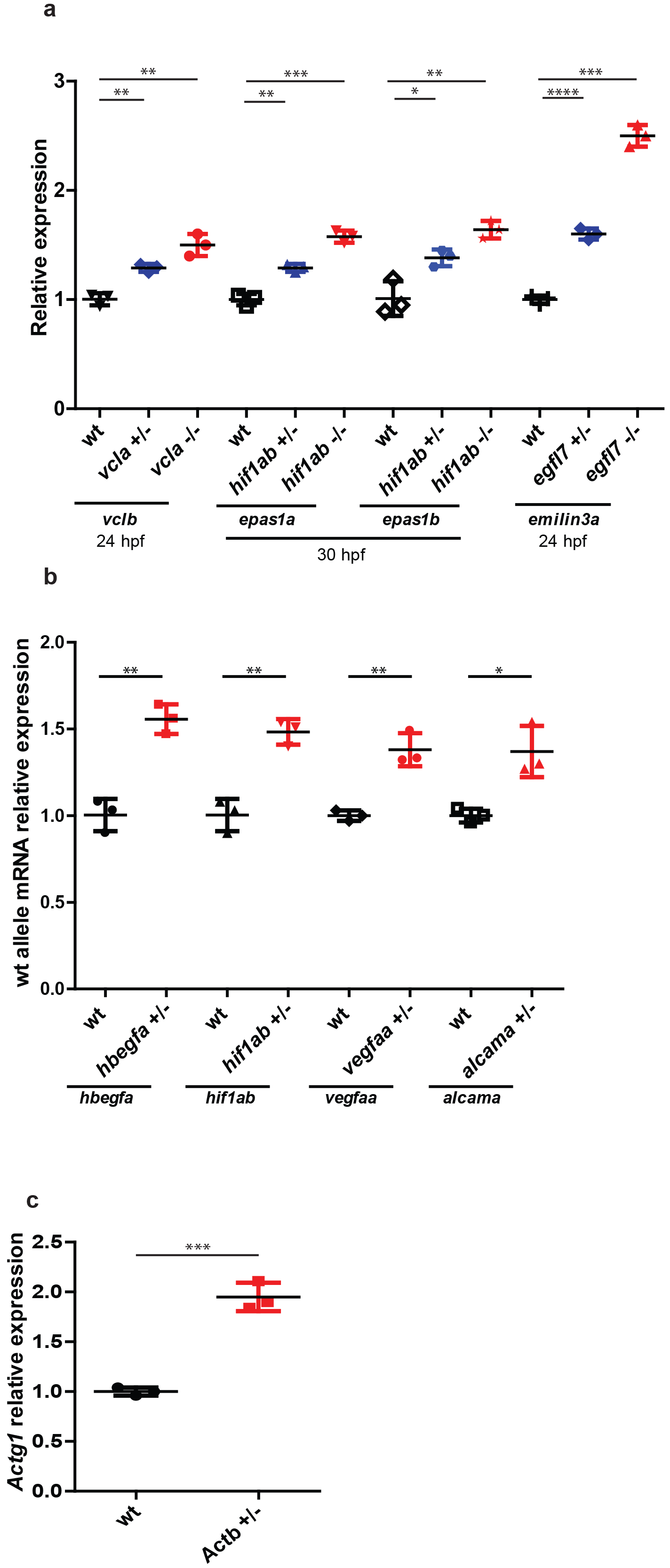

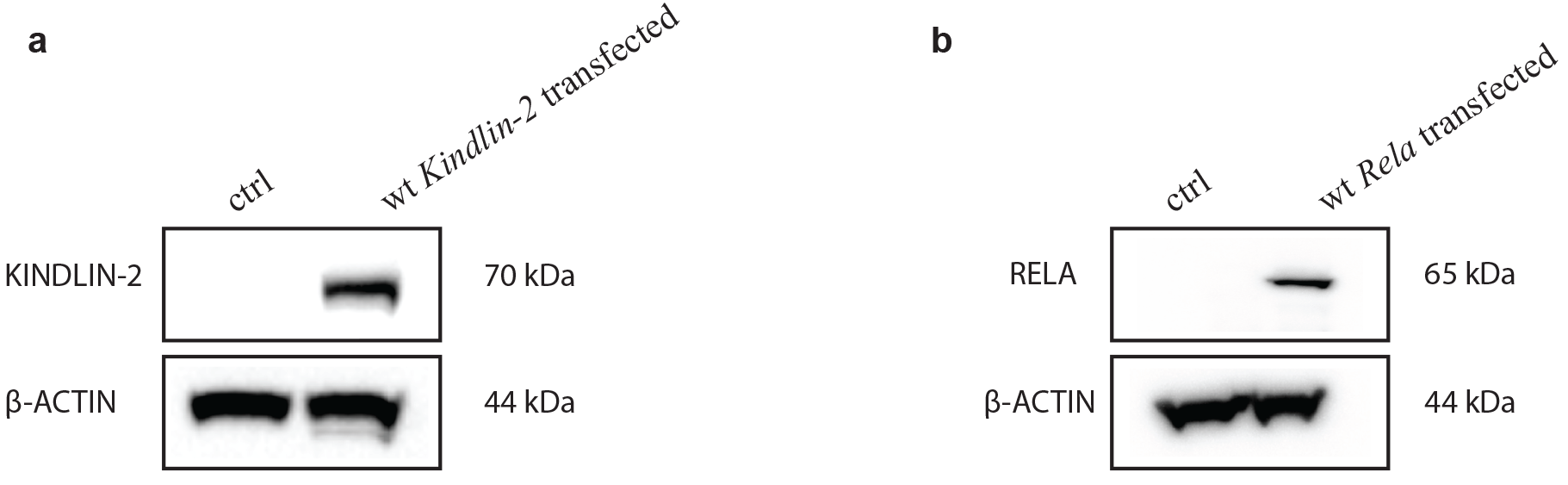

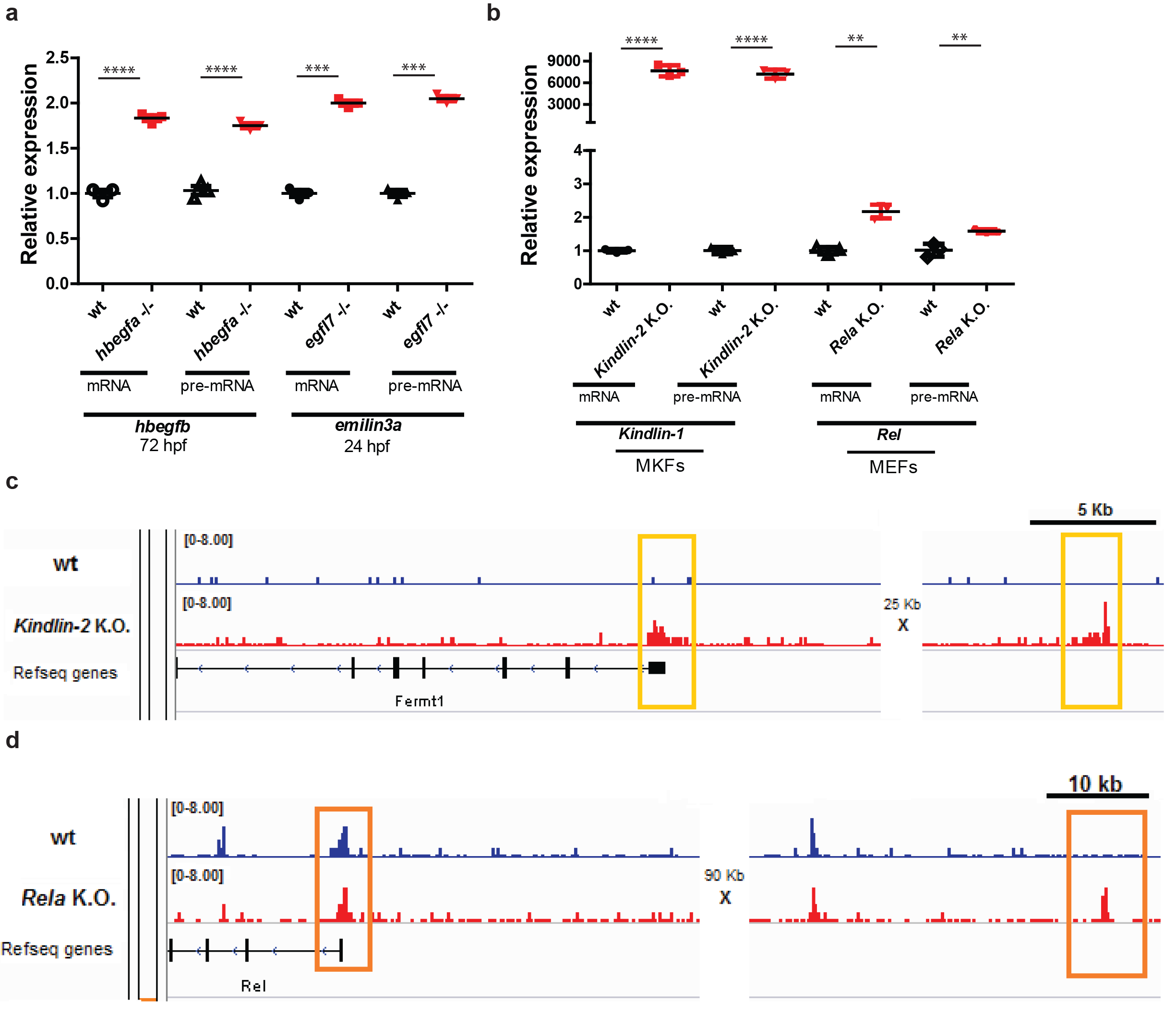

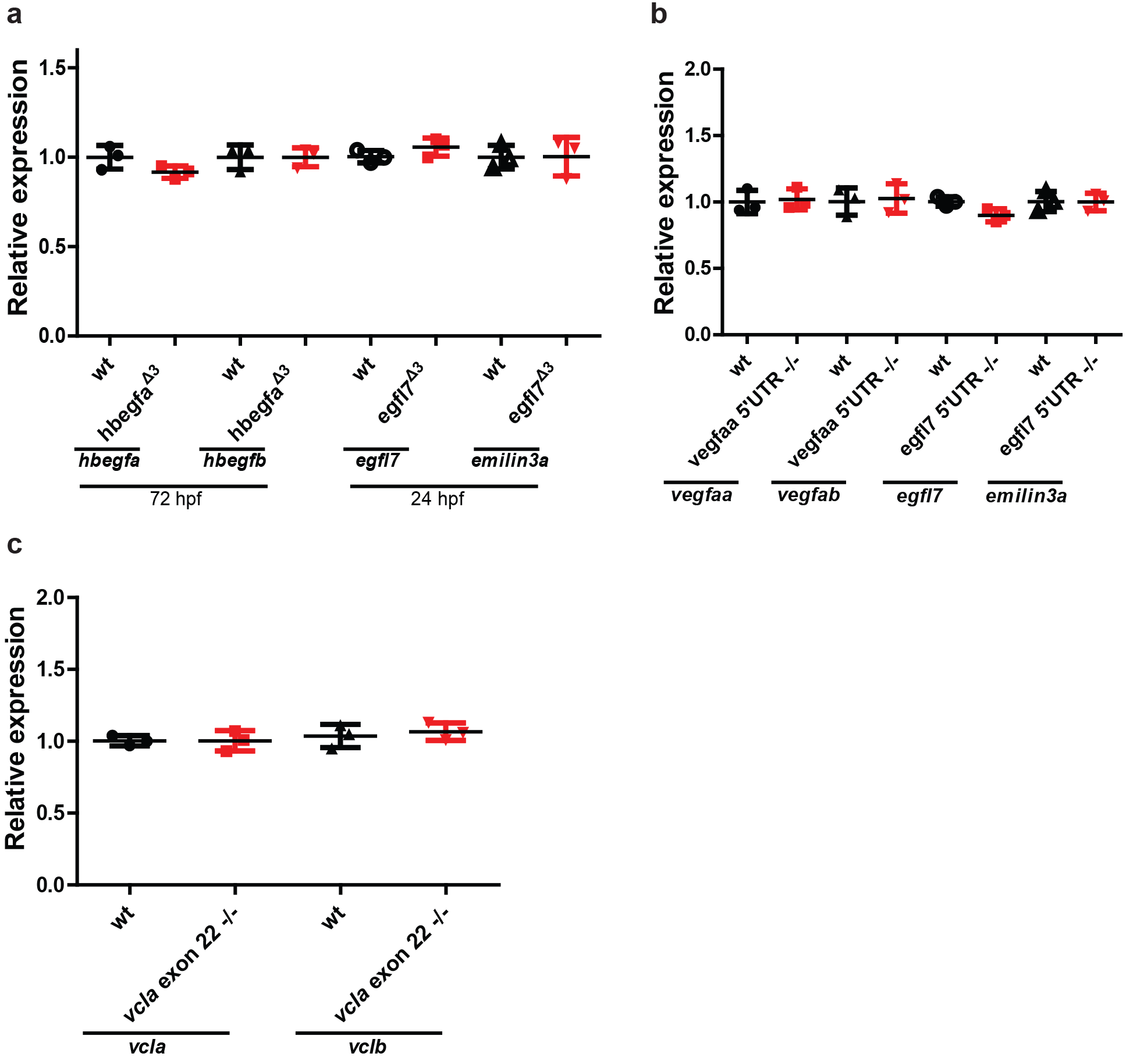

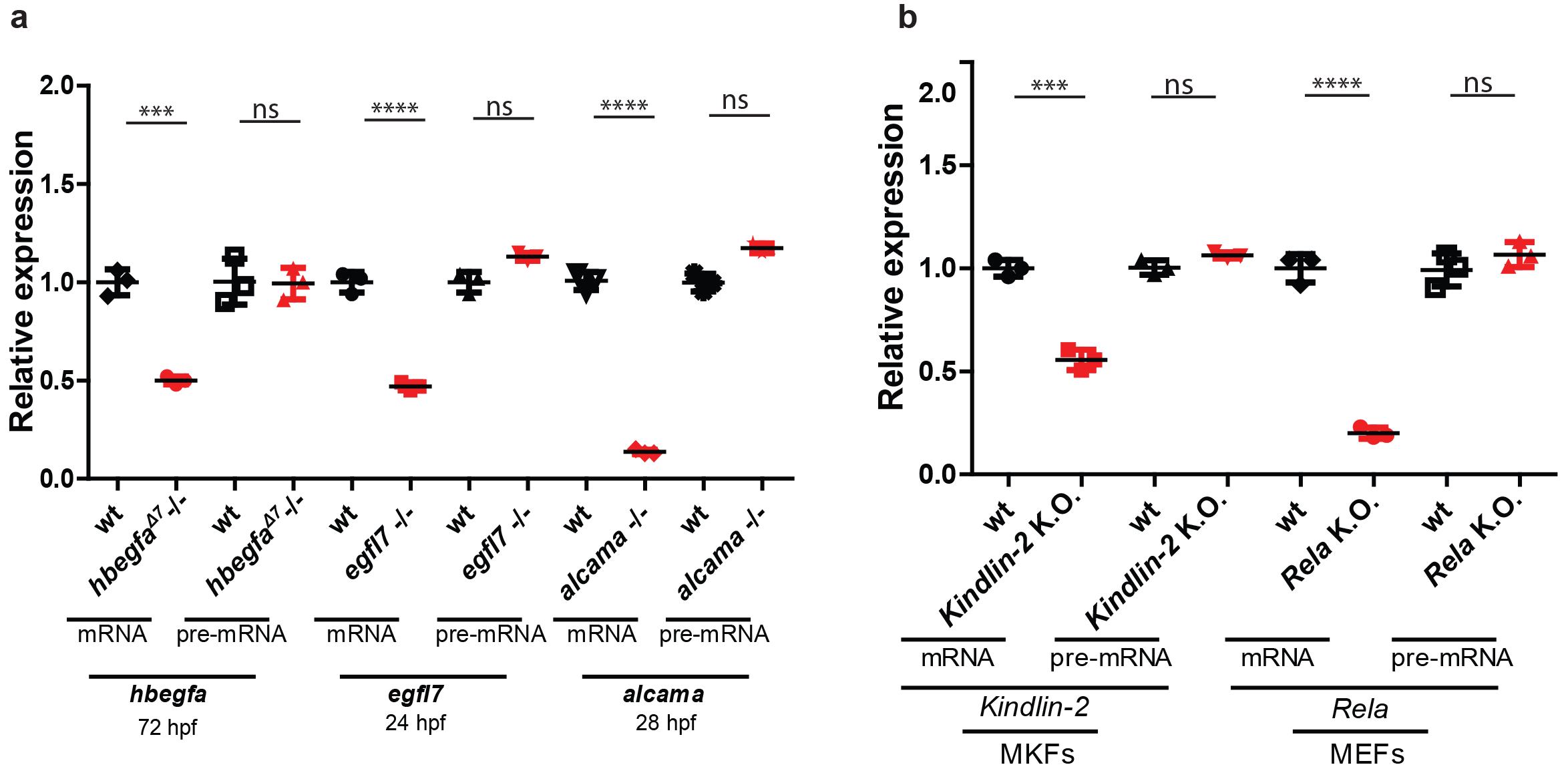

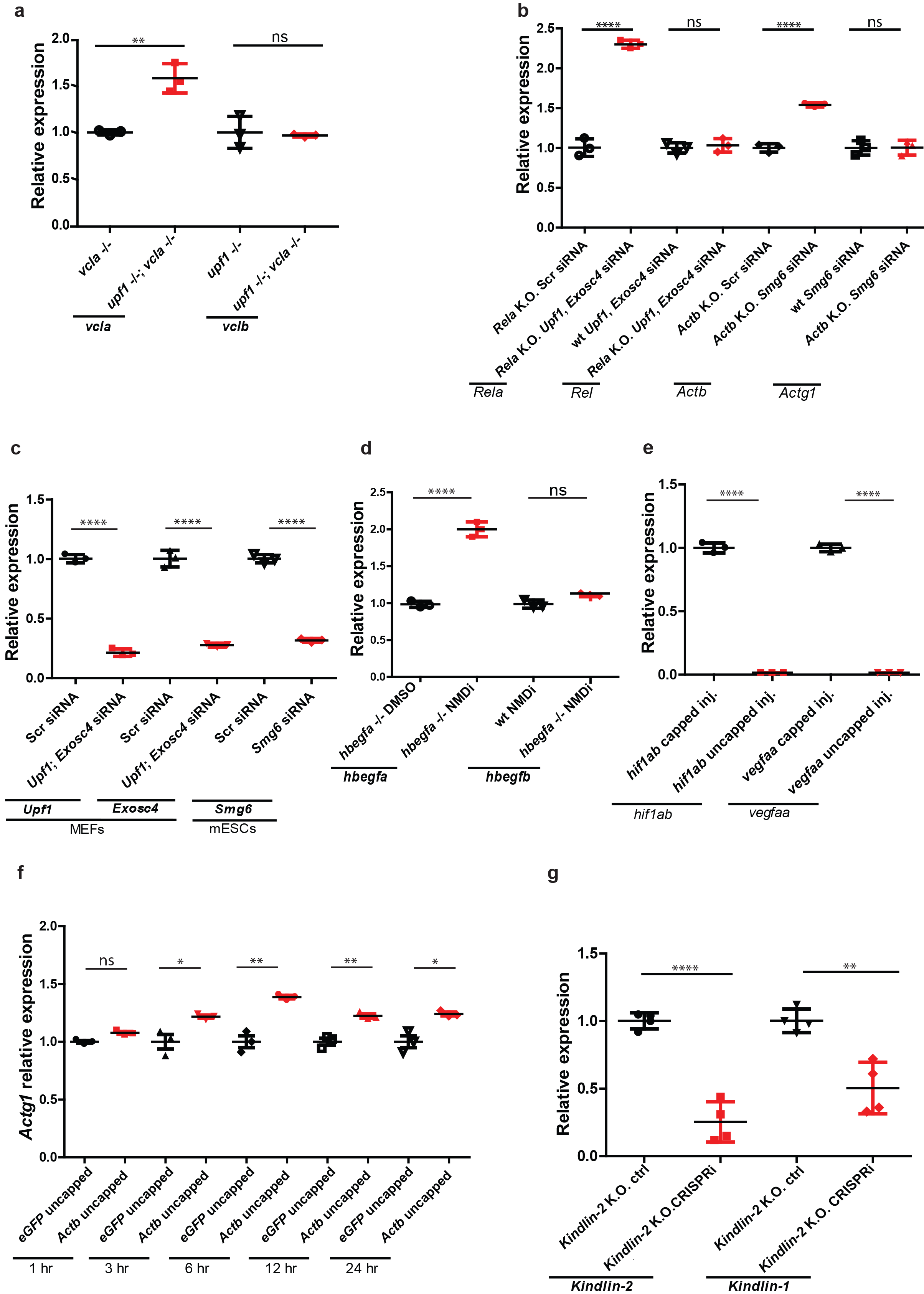

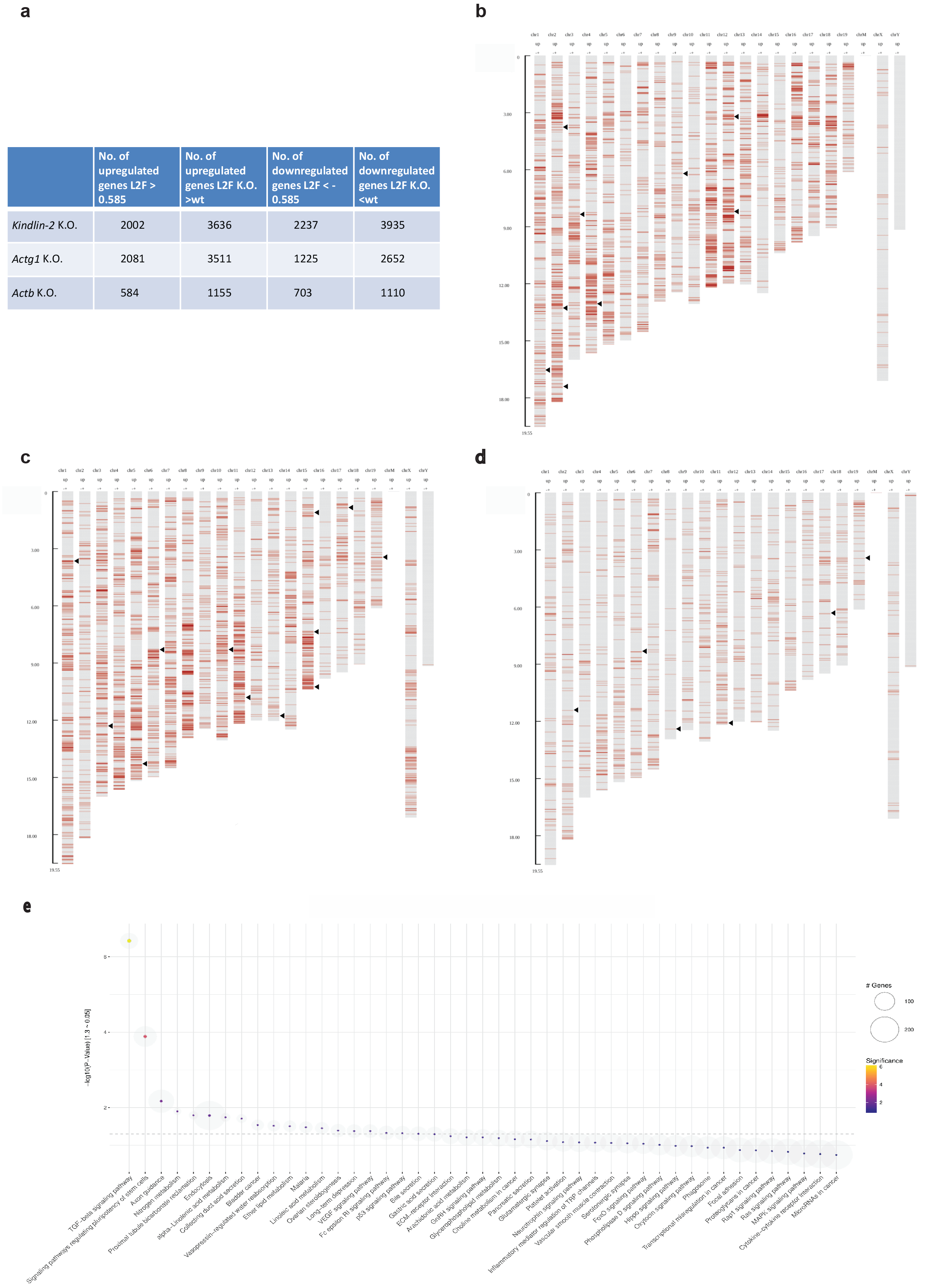

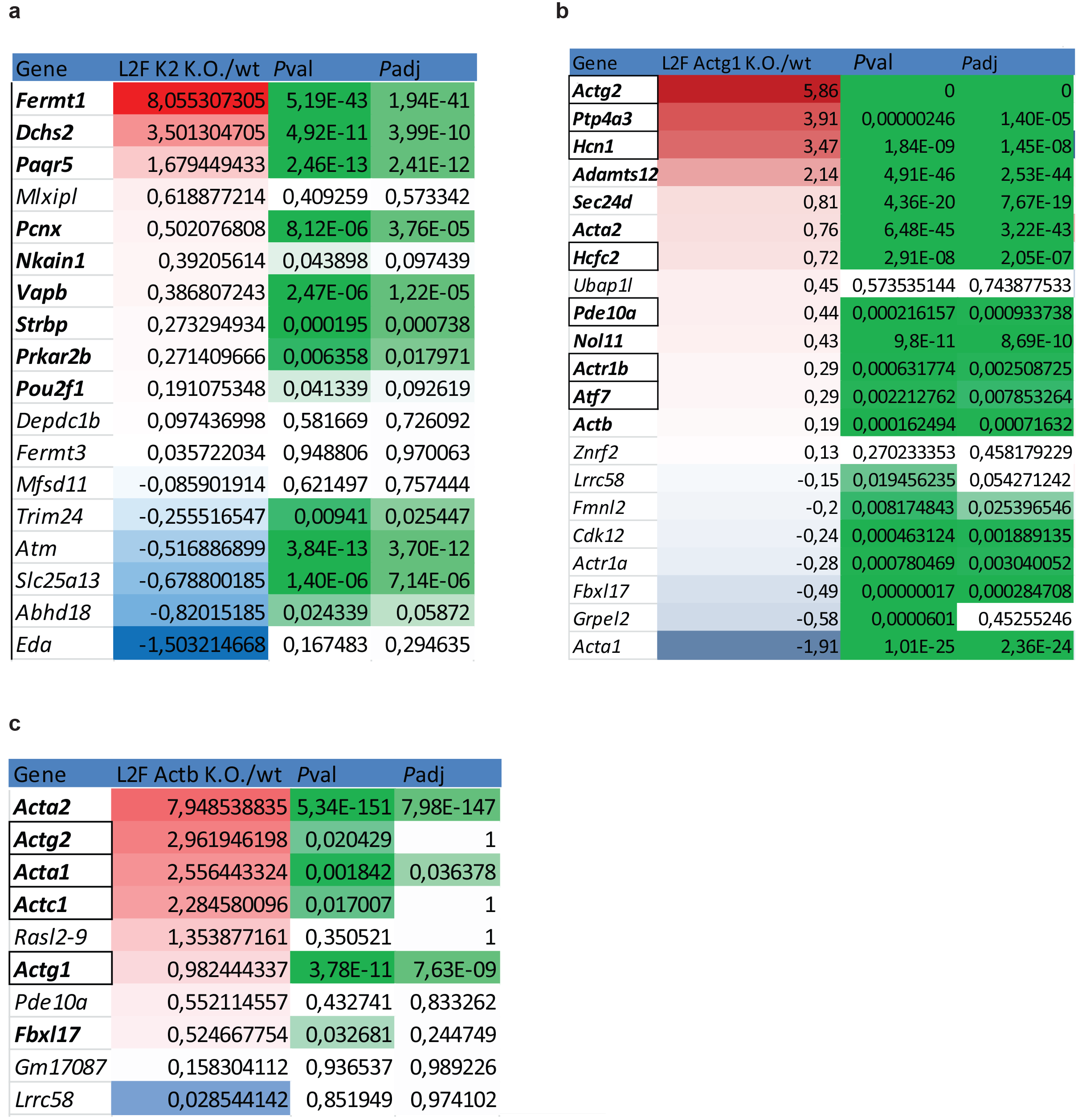

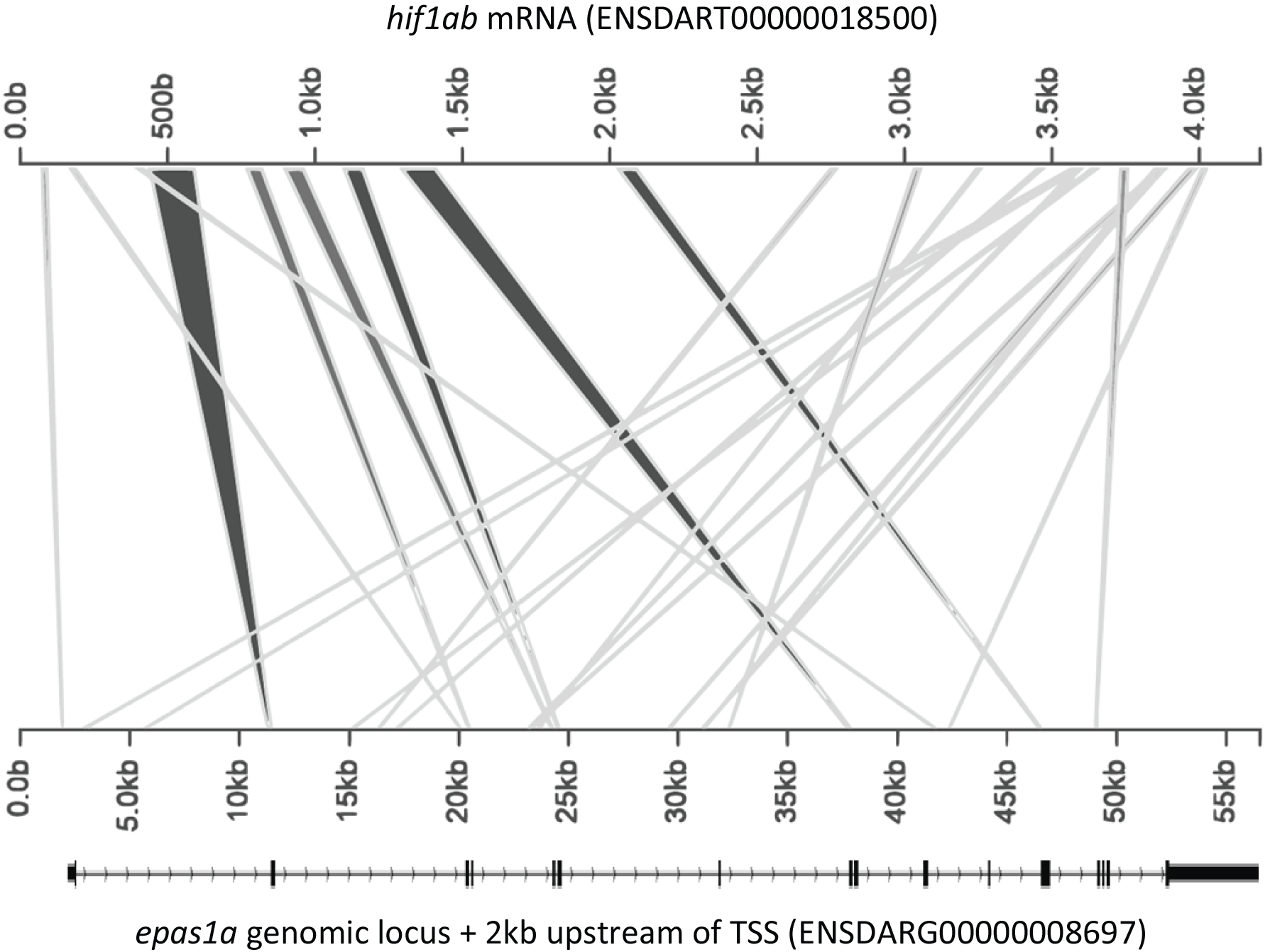

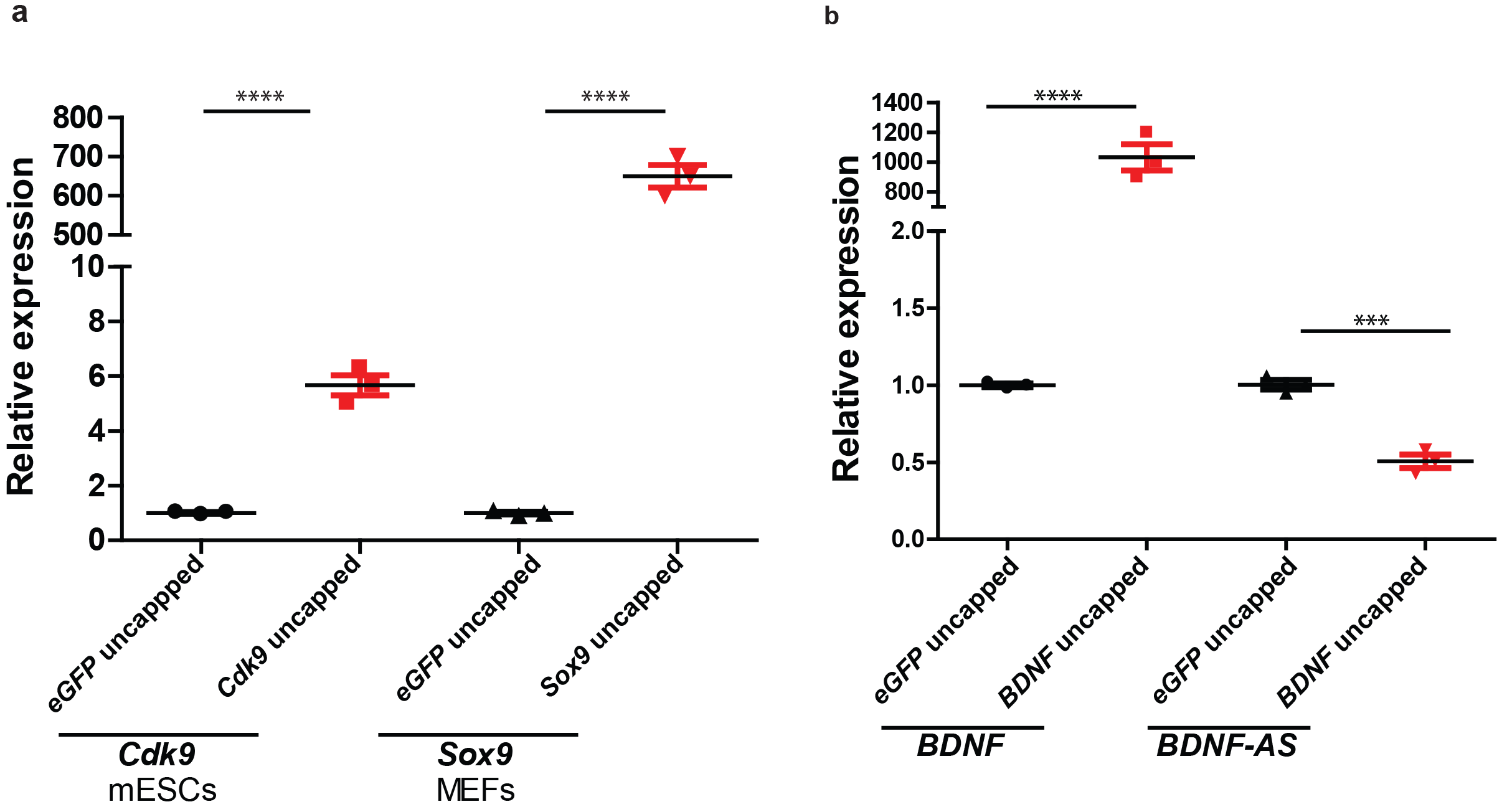

